# Extracellular loop 2 of G protein-coupled olfactory receptors is critical for odorant recognition

**DOI:** 10.1101/2021.10.26.465980

**Authors:** Yiqun Yu, Jody Pacalon, Zhenjie Ma, Lun Xu, Christine Belloir, Jeremie Topin, Loïc Briand, Jérôme Golebiowski, Xiaojing Cong

**Author notes:** Equal contribution (Yiqun Yu, Jody Pacalon, Zhenjie Ma).

## Abstract

G protein-coupled olfactory receptors (ORs) enable us to detect innumerous odorants. They are also ectopically expressed, emerging as attractive drug targets. ORs can be promiscuous or highly specific, which is part of Nature’s strategy for odor discrimination. This work demonstrates that the extracellular loop 2 (ECL2) plays critical roles in OR promiscuity and specificity. Using site-directed mutagenesis and molecular modeling, we constructed 3D OR models in which ECL2 forms a lid of the orthosteric pocket. ECL2 controls the shape and the volume of the odorant-binding pocket, maintains the pocket hydrophobicity and acts as a gatekeeper of odorant binding. The interplay between the specific orthosteric pocket and the variable, less specific ECL2 controls OR specificity and promiscuity. The 3D models enabled virtual screening of new OR agonists and antagonists, exhibiting 78% hit rate in cell assays. This approach can be generalized to structure-based ligand screening for other GPCRs that lack high-resolution 3D structures.

## Introduction

G protein coupled receptors (GPCRs) are the largest family of membrane proteins in the human genome, comprising over 800 members, half of which are olfactory receptors (ORs) ^1^. GPCRs detect diverse ligands and control most of the cell signaling. Despite their diverse functions, GPCRs conserve a seven transmembrane helical architecture (TM1–TM7), connected by 3 extracellular and 3 intracellular loops (ECL1–ECL3 and ICL1–ICL3). ORs belong to class A GPCRs which account for ~85% of the human GPCR genes. The orthosteric ligand-binding pocket in class A GPCRs is located within the extracellular half of the TM bundle, extending ~15 Å deep into the cell membrane ^2^. The pocket may be solvent accessible (e.g. in receptors for peptides or soluble molecules) or shielded by ECL2 (e.g. in lipid receptors and rhodopsin) ^3^. ECL2 is often the longest extracellular loop, which is highly variable in length, sequence and structure ^4–5^. A disulfide bond between ECL2 and TM3 is conserved in 92% of human GPCRs ^6^. It is important for ligand-binding and receptor activation ^3^. Peptide-activated GPCRs mostly contain an ECL2 in the form of a β-hairpin lying on the rim of the orthosteric pocket. ECL2 of GPCRs that are modulated by small-molecule endogenous ligands exhibits diverse shapes. They are often unstructured and cover partially or fully the pocket entrance ^3^. Rhodopsin is a case in-between, in which a β-hairpin shaped ECL2 inserts deep into the orthosteric pocket ^7^. It has been suggested that rhodopsin ECL2 represents an evolutionary transition between peptide receptors and small molecule receptors ^5^. In small molecule receptors, ECL2 may have evolved to mimic the peptide ligands and occupy part of the pocket, which renders the pocket suitable for binding small molecules. ECL2 plays important roles in ligand binding and activation of class A GPCRs ^4^. It may act as a gateway to the orthosteric pocket ^8–12^, bind allosteric modulators ^13–14^ or participate in receptor activation ^15–16^.

ECL2 of ORs are among the longest in class A GPCRs. ORs can be promiscuous or highly specific, in which ECL2 may play a central role. However, the lack of high-resolution OR structures hampers the study of OR-odorant recognition. In this work, we studied the role of ECL2 in two prototypical mouse ORs of the same subfamily, mOR256-3 and mOR256-8, which share 54% sequence identity. Our previous work indicated that mOR256-3 is promiscuous for a series of commonly encountered odorants, whereas mOR256-8 is rather specific ^17^. In this study, we found that ECL2 properties strongly modulate OR-odorant recognition. We performed site-directed mutagenesis along ECL2 and built 3D OR models that comply with the mutagenesis data. Virtual screening using the 3D models identified new mOR256-3 ligands, including an antagonist which selectively inhibited some of the agonists. The 3D models provide structural explanations to the promiscuity of mOR256-3 and the selective antagonism.

## Results

### Sequence analysis of OR ECL2

Sequence alignment of 1521 human and mouse ORs showed that their ECL2 mostlt contain 34-35 amino acids (Fig. 1). They are longer than ECL2 in most class A GPCRs. Three cysteines are highly conserved (C169, C179 and C189 in mOR256-3, conserved in 93.4%, 99.5% and 95.0% of human and mouse ORs, respectively). While C179 forms the classic disulfide bond with TM3, C169 and C189 have been suggested to form a second disulfide bond within ECL2 ^18^. Residues around the two disulfide bonds show high conservation, whereas the rest of the OR ECL2 sequence has diversified intensively (Fig. 1). It is plausible that the two disulfide bonds are important for ECL2 structuration and OR functions.

**Figure 1.**
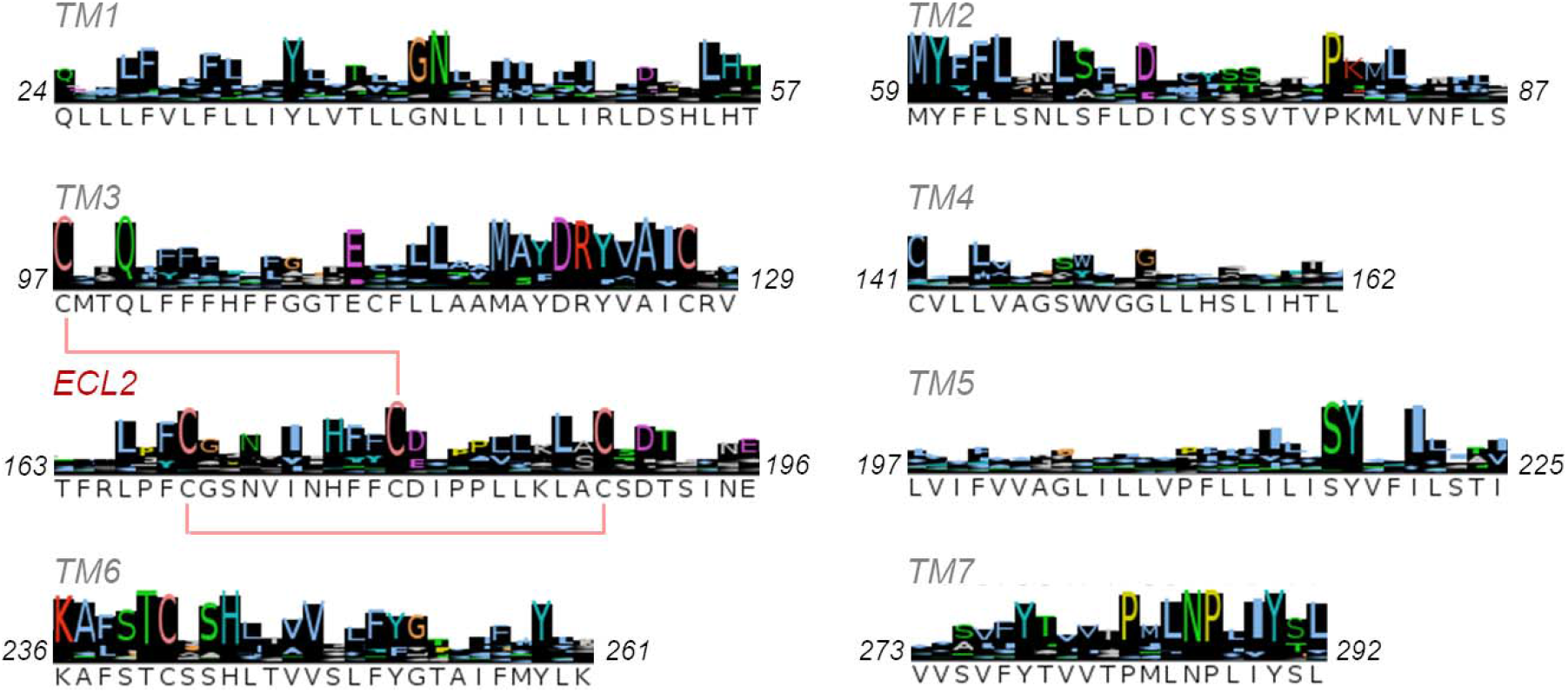
Consensus sequence of the TM regions and ECL2 in human and mouse ORs. Residue numbers in mOR256-3 are labeled on both sides of each region. Histogram indicates sequence conservation.

### Non-specific roles of ECL2 in OR responses to odorants

In our previous work, we screened diverse odorants at a near-saturating concentration (300 μM) on several ORs in the heterologous Hana3A cells. We found a wide range of potential ligands for mOR256-3 but only two for mOR256-8 ^17^. Yet, one or few point mutations in mOR256-8 could significantly expand its ligand spectrum ^17^. Here, we reexamined 20 of these odorants at various concentrations in heterologous cells expressing mOR256-3 or mOR256-8. Ten odorants activated mOR256-3 in a dose-dependent manner, including cyclic and acyclic alcohols, aldehydes, acids, ketones, and esters: R-carvone, coumarin, 1-octanol, allyl phenylacetate, benzyl acetate, citral, geraniol, 2-heptanone, octanal, and octanoic acid (Table S1 and Fig. S1A). mOR256-8 responded only to 1-octanol and geraniol in a dose-response manner, which are two primary acyclic alcohols of similar lengths (Table S1 and Fig. S1B).

Focusing on the role of ECL2, we performed site-directed mutagenesis to probe the residues that are responsible for the functional differences between mOR256-3 and mOR256-8. Based on the 3D models in our previous work ^17, 19–21^, we mutated 14 residues on TM3–TM6 around the orthosteric pocket, as well as 15 residues in ECL2 of mOR256-8 that differ from mOR256-3. In the narrowly-tuned mOR256-8, these residues were mutated one by one into their counterpart in the broadly-tuned mOR256-3. We then tested the response of the mutant receptors to R-carvone and coumarin, two reference ligands of mOR256-3. While wild-type (wt) mOR256-8 does not respond to these odorants, 14 of the mutants showed dose-dependent responses to R-carvone, and 9 of them also responded to coumarin (Fig. 2 A and B). Four of the mutations were in ECL2 (R173I, N175D, L181V, and L184M), which led to responses to both odorants (Fig. 2B). These residues flank the ECL2-TM3 disulfide bond, suggesting that this region (residues 173–184) is important for the receptor function. Five residues in this region are conserved in mOR256-8 and mOR256-3 (H176, F177, E180, P182, and A183). Therefore, we mutated the five residues in mOR256-3, in order to evaluate their role in this promiscuous receptor. They were mutated into alanine, except for A183 which was mutated into a bulky isoleucine. All the five mutations in mOR256-3 ECL2 systematically diminished the receptor’s response to R-carvone, coumarin and geraniol (Fig. 2C).

**Figure 2.**
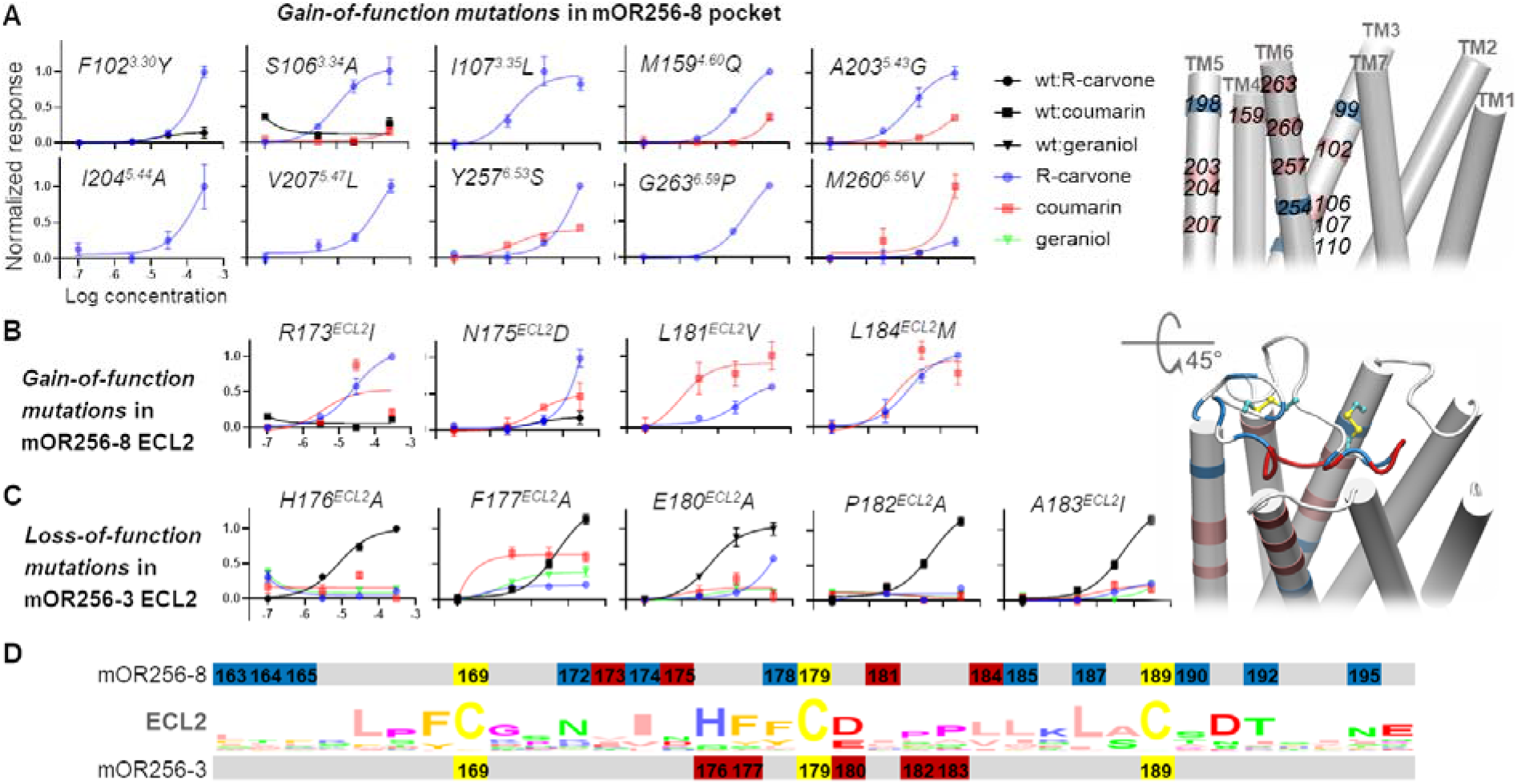
Site-directed mutagenesis and location of the mutation sites. Mutations in **(A)** mOR256-8 orthosteric pocket, **(B)** mOR256-8 ECL2 and (**C**) mOR256-8 ECL2 affected the response to various odorants. (**D**) Consensus ECL2 sequence and location of the mutation sites. Effective mutations are colored in pink (in A) and red (in D). Non-effective mutations are colored in blue, including V99^3.27^A, V110^3.38^T, L198^5.38^E, S254^6.50^T, R172^ECL2^N, I174^ECL2^L, L178^ECL2^F, I185^ECL2^L, M187^ECL2^L, V190^ECL2^T, A192^ECL2^T, and V195^ECL2^N in mOR256-8. In the 3D models, consistently, the non-effective mutation sites (blue) do no constitute the ligand binding site or the pathway to the binding site.

We also generated a chimeric mOR256-8 in which ECL2 was replaced with that of mOR256-3. However, it did not gain response to the ligands of mOR256-3. The above data highlight that residues 173–184 in ECL2 are critical but not solely responsible for ligand recognition or receptor promiscuity. Interestingly, while the mutations in the pocket affected the specific response to one ligand, mutations of residues 173–184 affected all the tested ligands in the same manner (gain or loss of response). Thus, the role of ECL2 in OR-ligand recognition appears less specific than the pocket. This is in line with the notion that in class A GPCRs, ECL2 acts as a vestibule or a molecular sieve of ligand binding and/or an allosteric site of receptor activation. Since residues 173–184 in ORs surround the conserved ECL2-TM3 disulfide bond, they are likely important in most, if not all, mammalian ORs. For instance, mutations in this region have dramatic impact on the response of mOR-EG to its odorants ^22^. This region has also been found to interact with the orthosteric ligands in several non-olfactory class A GPCRs ^4^.

### 3D modeling explains OR promiscuity

To date, there are no high-resolution OR structures or structural information on the structural fold of OR ECL2. We used circular dichroism (CD) spectroscopy of a truncated mOR256-3 ECL2 peptide to estimate its secondary structure content. The deconvolution of the CD spectrum revealed a largely unstructured peptide composed of 30% β-sheet, 13% helix, 6% turns and 51% random coil (Fig. S2). Nevertheless, the structure may be significantly different in a full-length OR, in which ECL2 is attached to TM3 by a disulfide bond. Therefore, we generated three types of 3D models using AlphaFold 2 ^23^, Modeller (A. Sali and T. L. Blundell, 1993), and Swiss-model ^24^. The three models displayed distinct structures in ECL2 (Fig. 3A and S3). We evaluated the quality of the models using site-directed mutagenesis data and docking. The model that best matched these data was generated by Modeller based on our hand-curated multiple sequence alignment (Fig. S4). In this model, ECL2 appears as an unstructured coil, in which residues 173–184 form a lid of the orthosteric pocket (Fig. 3A). While residue 181 is ~8 Å directly above the “toggle switch residue” Y^6.48^, residues 180–183 may interact directly with the ligands (Fig. 3A). The model also suggests that the pocket of mOR256-3 is much larger than mOR256-8, showing two connected cavities (Fig. 3B). This may allow mOR256-3 to accommodate odorants of diverse size and shape. Molecular docking suggests that the upper cavity prefers the cyclic ligands whereas the deeper cavity accommodates the acyclic ones (Fig. 3B). The pocket of mOR256-8 shows only one small cavity for its acyclic ligands. We estimated the pocket volume of all the human and mouse ORs by simply summing up the side-chain volume of the residues outlining the pocket with or without ECL2. We found that the pocket of mOR256-3 is of an average size whereas that of mOR256-8 is among the smallest (Fig. S5). Thus, the larger pocket volume of mOR256-3 than mOR256-8 may provide a structural explanation to the promiscuity of the former. To assess this hypothesis and the model quality, we use the model to virtually screen for new mOR256-3 ligands by molecular docking.

**Figure 3.**
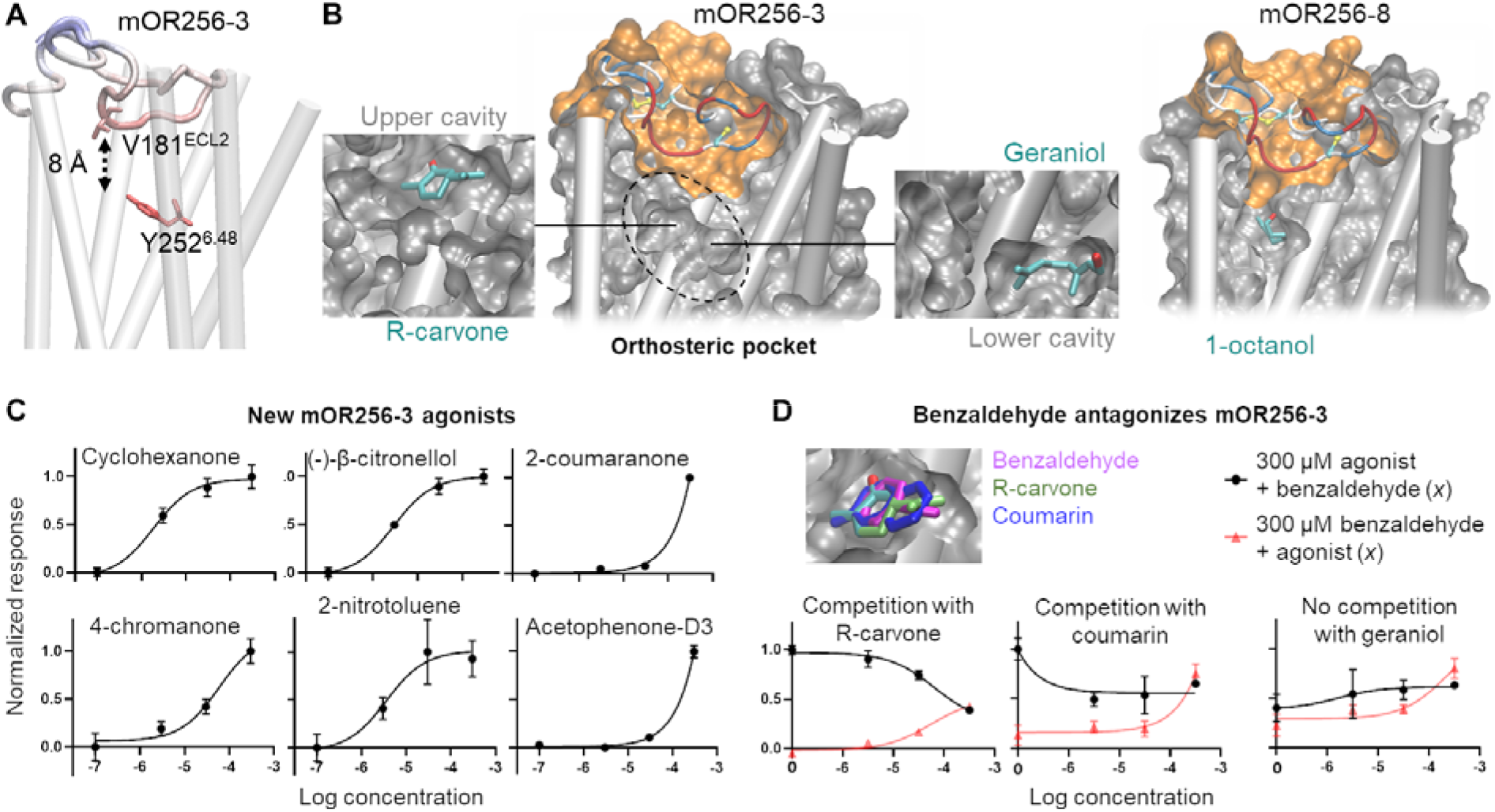
Selected 3D models and new mOR256-3 ligands discovered by virtual screening. (**A**) mOR256-3 model that best match the mutagenesis data. ECL2 is colored by the distance to the toggle switch Y^6.48^. (**B**) Cross section of the best model of mOR256-3 and mOR256-8, illustrating ECL2 as the pocket lid. mOR256-3 displays two connected cavities in the pocket, in which the upper cavity binds cyclic ligands and the lower one accommodates acyclic molecules. **(C)** Dose-dependent curves of new mOR256-3 agonists from virtual screening. (**D**) Benzaldehyde binds in the same cavity as R-carvone and coumarin. It inhibits R-carvone (partially) and coumarin (slightly) but not geraniol.

Docking benchmarks were first performed 54 compounds, including 10 known ligands of mOR256-3 and 42 decoys (Table S2) ^17^. An ensemble-docking protocol (Fig. S6) was used to account for the conformational flexibility of the OR. Briefly, molecular dynamics (MD) simulations were performed on the initial model of mOR256-3 to sample the receptor conformations. From the MD trajectory, 20 snapshots were extracted for ensemble docking using Autodock Vina ^25^. The in-house model–generated by Modeller and selected according to site-directed mutagenesis data–gave the best predictions on the benchmark compounds compared to those generated by AlphaFold2 and the Swiss model (Table 1). Removing ECL2 from this model significantly reduced the model’s predictivity (Table 1).

**Table 1.** Docking benchmark using different 3D models of mOR256-3 and 54 compounds.

| Model | MD snapshot<sup>a</sup> | MCC | Hit rate (precision) | Recall | True positive | True negative | False positive | False negative |
| --- | --- | --- | --- | --- | --- | --- | --- | --- |
| In-house model | 1 | 0.50 | 0.60 | 0.60 | 6 | 38 | 4 | 4 |
|  | 2 | 0.26 | 0.40 | 0.40 | 4 | 36 | 6 | 6 |
| In-house model without ECL2 | 1 | 0.26 | 0.40 | 0.40 | 4 | 36 | 6 | 6 |
|  | 2 | 0.13 | 0.30 | 0.30 | 3 | 35 | 7 | 7 |
| Swiss model | 1 | 0.23 | 0.36 | 0.40 | 4 | 35 | 7 | 6 |
|  | 2 | 0.20 | 0.33 | 0.40 | 4 | 34 | 8 | 6 |
| AlphaFold2 | 1 | 0.38 | 0.50 | 0.50 | 5 | 37 | 5 | 5 |
|  | 2 | 0.26 | 0.40 | 0.40 | 4 | 36 | 6 | 6 |

<sup>a</sup> Two snapshots that gave the best MCC.

We then chose the two best performing snapshots of the above model to virtually screen an in-house library of 80 odorants. The screening returned 9 hits, which were tested in functional assays in Hana3A cells. Six of them turned out to be mOR256-3 agonists and one (benzaldehyde) was a partial antagonist, giving 78% hit rate (Fig. 3C-D and Table S3). Benzaldehyde antagonized R-carvone and coumarin but not geraniol (Fig. 3D). This is consistent with the model prediction that benzaldehyde binds in the same sub-cavity of the mOR256-3 pocket as R-carvone and coumarin, whereas geraniol binds in the other cavity (Fig. 3D). These findings indicate that OR promiscuity results from the combination between pocket volume and ECL2 conformation.

### ECL2 controls pocket shape and hydrophobicity

To further examine the role of ECL2 in odorant recognition, we constructed three mOR256-3 chimeras, by replacing its ECL2 with that of M2 muscarinic receptor, β2 adrenergic receptor and 5HT serotonin 2C receptor, respectively (denotated as ch-β_2_AR^ECL2^, ch-M_2_R^ECL2^ and ch-5HT_2C_R^ECL2^). ECL2 of these receptors exhibit distinct structures (Fig. 4A). In Hana3A cells, the chimeras showed no significant response to the mOR256-3 ligands. Nevertheless, they all displayed specific dose-dependent response to trans-cinnamaldehyde (Fig. 4B), while wt mOR256-3 does not respond to this odorant ^17^. To understand how the chimeric mOR256-3 became specific receptors of trans-cinnamaldehyde, we built homology models for the chimeras and performed all-atom molecular dynamics (MD) simulations in an explicit membrane-water environment. The homology models were built by assuming that ECL2 of the chimeras preserve their native fold in β_2_AR, M_2_R and 5HT_2C_R, respectively. The models illustrated that ECL2 of the chimeras only partly covered the ligand entrance. Unlike wt mOR256-3 in which ECL2 keeps the orthosteric pocket hydrophobic, the pocket of the chimeras was hydrated during the MD (Fig. 4A). This might be the reason why the chimeras did not respond to the hydrophobic ligands of mOR256-3. Rather, they prefered the less hydrophobic trans-cinnamaldehyde (Table S1).

**Figure 4.**
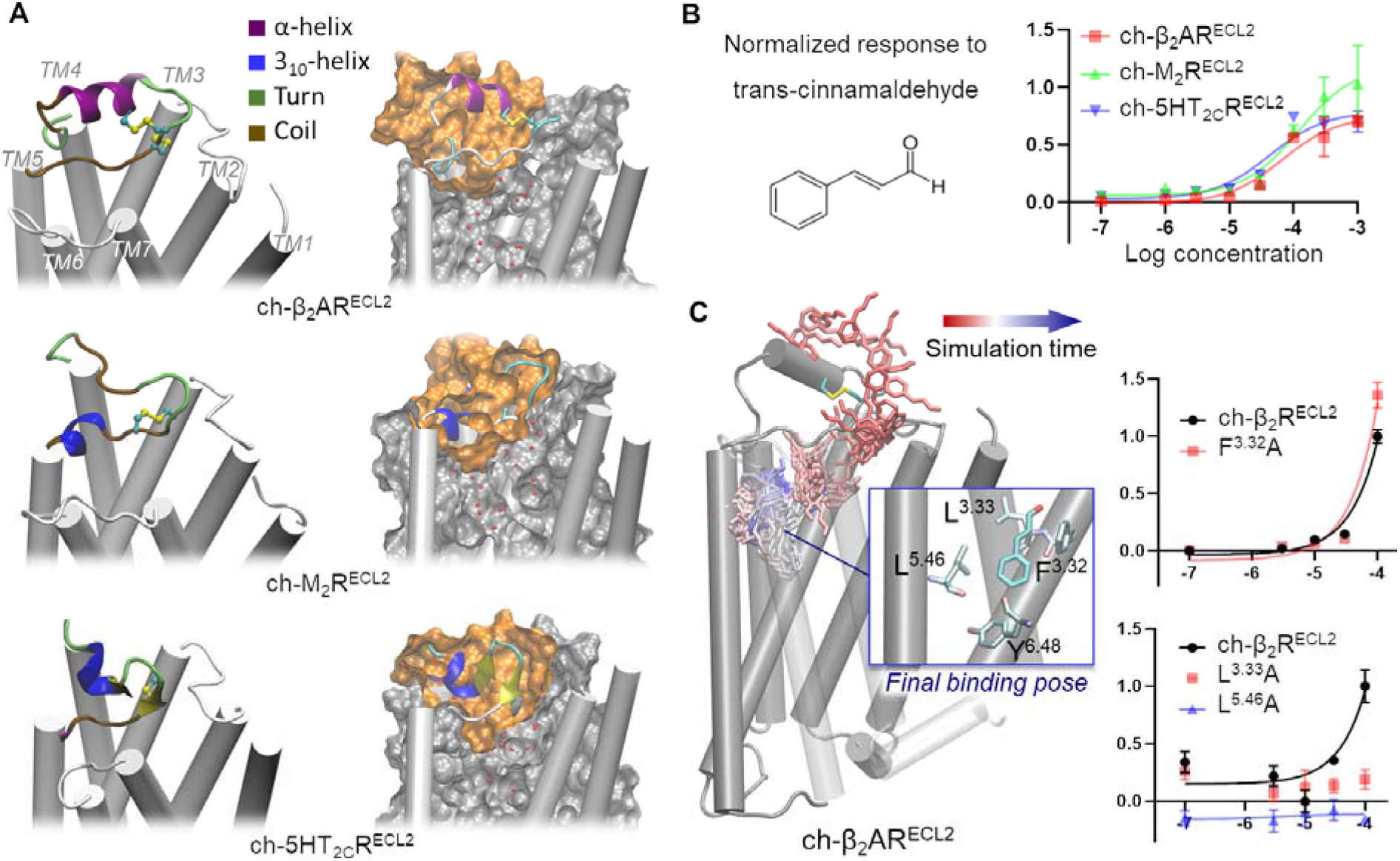
Structures and function of mOR256-3 chimeras. (**A**) Homology models of mOR256-3 chimeras with different ECL2 sequences and structures. MD simulations illustrate that the pocket of the chimeras is hydrated. (**B**) Dosedependent responses of the three chimeras to trans-cinnamaldehyde. (**C**) Trans-cinnamaldehyde entered the pocket of β_2_AR^ECL2^ via the ECL2-TM7 gap and stabilized in a binding pose that activates the toggle switch Y^6.48^. Mutating the trans-cinnamaldehyde-binding residues affect the receptor response to this ligand.

We then added trans-cinnamaldehyde in the MD simulations of wt mOR256-3 and the chimeras to monitor the ligand binding. The ligand was initially placed 10 Å above ECL2 and was restrained within a 15 Å radius around ECL2. Each system underwent 30 independent MD runs of 200 ns. Only two binding events occurred, both in ch-β_2_AR^ECL2^, in which trans-cinnamaldehyde could enter deep in the orthosteric pocket and interact with the toggle switch residue Y6.48 (Fig. 4C). It caused the side chain of Y6.48 to flip toward TM5, which is likely an early step of OR activation ^20^. In the case of wt mOR256-3, trans-cinnamaldehyde associated with ECL2 but could not enter the pocket. The binding pose of trans-cinnamaldehyde in ch-β_2_AR^ECL2^ suggests that wt mOR256-3 cannot accommodate this ligand, since ECL2 occupies part of its pocket (Fig. 2A-B). Indeed, mOR256-3 ligands are generally smaller or more flexible than trans-cinnamaldehyde. To verify the binding pose of trans-cinnamaldehyde observed in the MD simulations, we mutated three pocket residues that are in close contact with the ligand. Mutations L^3.33^A and L^5.46^A abolished the receptor response to trans-cinnamaldehyde, whereas F^3.32^A slightly increased the response efficacy (Fig. 4C). The results suggest that the recognition of trans-cinnamaldehyde is specific to the orthosteric pocket, whereas ECL2 served as an unspecific molecular sieve for the ligand entrance.

## Discussion

Mammalian OR sequences have highly diversified during evolution to detect and discriminate a vast spectrum of odorants. Specific (or narrowly-turned) ORs may be responsible for the detection of specific odorants or endogenous ligands when ectopically expressed ^26–29^. Promiscuous (or broadly-tuned) ORs may play exert important functions in olfaction, such as expanding the detection spectrum, diversifying the combinatorial code, and acting as general odor detectors or odor intensity analyzers ^17^. We have previously found that promiscuous ORs feature mostly non-polar interactions in the orthosteric pocket with odorants, which are more adaptable to different odorant structures ^21^. Here, we showed that ECL2 is indispensable for OR promiscuity. ECL2 acts as a pocket lid to maintain the pocket hydrophobicity and also forms the upper part of the pocket to control its shape and volume. It interacts directly with the odorants, while its structural flexibility and mostly hydrophobic nature may tolerate diverse odorants, resulting in promiscuity. Indeed, in class A GPCRs, ECL2 may change conformations upon ligand binding and adopt different forms for different ligands ^4^. The evolution of ECL2 in class A GPCRs is strongly coupled to that of the orthosteric pocket ^5^. Therefore, class A GPCR-ligand recognition relies on the interplay between ECL2 and the orthosteric pocket. ECL2 may also take part in receptor activation via allosteric coupling with the receptor movements on the intracellular side ^4^. However, this aspect is beyond the scope of the current study. Note that the 3D models reported here are not to present the exact structural fold of ECL2. Rather, they are to illustrate the approximate position of the ECL2 residues according to the mutagenesis data, as well as to serve virtual screening for new odorants. The screening performance indicates that such models are suitable structural basis for ligand discovery. The virtual screening approach used here may serve the design of biosensors with wide odor detection spectrum, or specific odor maskers and/or drug candidates targeting ectopic ORs.

## Materials and Methods

### Chemicals and OR constructs

Odorants were purchased from Sigma Aldrich. They were dissolved in DMSO to make stock solutions at 1 mM then diluted freshly in optimal MEM (ThermoFisher) to prepare the odorant stimuli. The OR constructs were kindly provided by Dr. Hiroaki Matsunami (Duke University, Durham, USA). Site-directed mutants were constructed using the Quikchange site-directed mutagenesis kit (Agilent Technologies). The sequences of all plasmid constructs were verified by both forward and reverse sequencing (Sangon Biotech, Shanghai, China).

### Chimera Construction

All chimeras were constructed by three PCR steps with modification ^30^. Briefly, two fragments were amplified from the mOR256-3 while ECL2 of β_2_AR, M_2_R and 5HT_2C_R was synthesized by Sangon Biotech Co. (Shanghai, China). The primers were partially complementary at their 5’ ends to the adjacent fragments, necessary to fuse the different fragments together. Three fragments were purified and fused together in a second PCR step. Equal amount of each fragment was mixed with dNTP and Phusion^®^ High-Fidelity DNA Polymerase (NEB) in the absence of primers. The PCR program consisted of 10 repetitive cycles with a denaturation step at 98 °C for 10 s, an annealing step at 55 °C for 30 s and an elongation step for 30 s at 72 °C. The third step corresponded to the PCR amplification of the fusion product using the primers of mOR256-3. The PCR product was purified and ligated into PCI vector. The sequences of all chimeras were verified by both forward and reverse sequencing.

### Cell culture and transfection

We used Hana3A cells, a HEK293T-derived cell line that stably expresses receptor-transporting proteins (RTP1L and RTP2), receptor expression-enhancing protein 1 (REEP1) and olfactory G protein (Gα_olf_) ^31^. The cells were grown in MEM (Corning) supplemented with 10% (vol/vol) fetal bovine serum (FBS; ThermoFisher) plus 100 μg/ml penicillin-streptomycin (ThermoFisher), 1.25 μg/ml amphotericin (Sigma Aldrich), and 1 μg/ml puromycin (Sigma Aldrich).

All constructs were transfected into the cells using Lipofectamine 2000 (ThermoFisher). Before the transfection, the cells were plated on 96-well plates (NEST) and incubated overnight in MEM with 10% FBS at 37 °C and 5% CO_2_. For each 96-well plate, 2.4 μg of pRL-SV40, 2.4 μg of CRE-Luc, 2.4 μg of mouse RTP1S, and 12 μg of receptor plasmid DNA were transfected. The cells were subjected to a luciferase assay 24 hours after transfection.

### Luciferase assay

The luciferase assay was performed with the Dual-Glo Luciferase Assay Kit (Promega) following the protocol in ref. ^31^. OR activation triggers the Gα_olf_-driven AC-cAMP-PKA signaling cascade and phosphorylates CREB. Activated CREB induces luciferase gene expression, which can be quantified luminometrically (measured here with a bioluminescence plate reader (MD SPECTRAMAX L)). Cells were co-transfected with firefly and *Renilla* luciferases where firefly luciferase served as the cAMP reporter. *Renilla* luciferase is driven by a constitutively active simian virus 40 (SV40) promoter (pRL-SV40; Promega), which served as a control for cell viability and transfection efficiency. The ratio between firefly luciferase versus *Renilla* luciferase was measured. Normalized OR activity was calculated as (*L*_N_ – *L*_min_)/(*L*_max_ – *L*_min_), where *L*_N_ is the luminescence in response to the odorant, and *L*_min_ and *L*_max_ are the minimum and maximum luminescence values on a plate, respectively. The assay was carried out as follows: 24h after transfection, medium was replaced with 100 μl of odorant solution (at different doses) diluted in Optimal MEM (ThermoFisher), and cells were further incubated for 4 h at 37 °C and 5% CO_2_. After incubation in lysis buffer for 15 minutes, 20 μl of Dual-Glo^™^ Luciferase Reagent was added to each well of 96-well plate and firefly luciferase luminescence was measured. Next, 20 μl Stop-Glo Luciferase Reagent was added to each well and *Renilla* luciferase luminescence was measured. Data analysis followed the published procedure in ref. ^31^. Three-parameter dose-response curves were fitted with GraphPad Prism 9.

### Molecular modeling

The in-house models of mOR256-3 and mOR256-8 were generated with Modeller 9.21 (A. Sali and T. L. Blundell, 1993) using our hand curated sequence alignment to four structure templates: human a2AR (pdb 2YDV), human CXCR1 (pdb 2LNL), human CXCR4 (pdb 3ODU) and bovin rhodospin (pdb 1U19). For each receptor, 2500 models were generated and evaluated by the helicity of their TM regions. The 250 top ranked models were selected and clustered using the k-means algorithm. We obtained 5 clusters for each receptor and selected a representative model that was the most compatible with the mutagenesis data. The swiss models were generated using the Swiss-model webserver ^24^ using default settings. The AlphaFold 2 models ^23^ were generated using the Södinglab API based on the MMseqs2 server (Mirdita et al., 2019) for the multiple sequence alignment creation. Docking was performed with Autodock Vina and default settings ^25^.

### MD simulations

The receptors or receptor-odorant complexes were embedded in a bilayer of POPC using PACKMOL-Memgen ^32^. Each system was solvated in a periodic 75 × 75 × 105 Å^3^ box of explicit water and neutralized with 0.15 M of Na^+^ and Cl^-^ ions. Effective point charges of the ligands were obtained by RESP fitting ^33^ of the electrostatic potentials calculated with the HF/6-31G* basis set using Gaussian 09 ^34^. The Amber 99SB-ildn ^35^, lipid 14 ^36^ and GAFF ^37^ force fields were used for the proteins, the lipids and the ligands, respectively. The TIP3P ^38^ and the Joung-Cheatham ^39^ models were used for the water and the ions, respectively.

The process of ligand binding was simulated with 30 runs of 200 ns of all-atom brute-force MD for each OR-ligand pair using Amber18. The ligand was initially placed 10 Å above ECL2. After energy minimization, each system was gradually heated to 310 K with a restraint of 200 kcal/mol on the receptor and the ligand, followed by 5 ns of pre-equilibration with a restraint of 5 kcal/mol, and 5 ns of unrestrained equilibration. During the production run, when the ligand exceeded 15 Å from the center of ECL2, a distance restraint of 10 kcal/mol was applied to drive the ligand toward the center.

To thoroughly sample the conformations of mOR256-3 for ensemble docking, we used an enhanced sampling technique, replica exchange with solute scaling (REST2) ^40^. REST2 MD was performed with 48 replicas in the *NVT* ensemble using Gromacs 5.1 ^41^ patched with the PLUMED 2.3 plugin ^42^. The protein and the ligands were considered as “solute” in the REST2 scheme-force constants of their van der Waals, electrostatic and dihedral terms were scaled down to facilitate conformational changes. The effective temperatures used for generating the REST2 scaling factors ranged from 310 K to 700 K, following a distribution calculated with the Patriksson-van der Spoel approach ^43^. Exchange between replicas was attempted every 1000 simulation steps. This setup resulted in an average exchange probability of ~40%. A total of 60 ns × 48 replicas of REST2 MD were carried out. The first 10 ns were discarded for equilibration and only the original unscaled replica (at 310 K effective temperature) was collected and analyzed.

### Circular dichroism

Peptides were chemically synthesized (Smart Bioscience SAS, France) according to the amino acid sequence of mOR256-3 ECL2. Cys179 was substituted with serine to avoid disulfide bonding with Cys169 or Cys189. The Cys169-Cys189 disulfide bond was correctly formed. Circular dichroism spectra of the peptides were measured on a Jasco J-815 spectropolarimeter equipped with a Peltier temperature control fixed at 20°C. Stock solutions of peptide were prepared at 10 mg/mL in 10 mM potassium phosphate buffer pH7, and further diluted in 10 mM potassium phosphate buffer, supplemented with trifluoroethanol (TFE) (1:1) at pH 5.5, which is known as a helix-enhancing cosolvent ^44^. Using a 0.01 cm path length quartz cell (Hellma), the peptide spectra were recorded between 178 to 260 nm. The spectra were corrected for buffer contributions and converted to mean residue ellipticity in deg cm^2^ dmol^-1^. The spectra were averaged over 10 scans accumulated at 1 nm intervals with 50-nm/min scan speed and 5 s response times. They were smoothed using the Stavitzky-Golay convolution filter with a span of 5. The secondary structure proportions were estimated using the deconvolution JWSSE-513 algorithm with Reed’s reference spectra, a protein structural analysis program suitable for peptide, included in Spectra Manager software (JASCO) ^45–46^.

## Author contributions

ZM, LX and WL performed the *in vitro* experiments; YY and XC analyzed the *in vitro* data; JP and XC performed the *in silico* modeling; JP, JT, XC and JG analyzed the *in silico* data; CB measured the CD spectra; CB and LB analyzed the CD data; JT contributed to the research design; XC, YY and JG designed the research and wrote the paper.

## Acknowledgements

This work was supported by the NeuroMod Institute of the University of Côte d’Azur (2019-2020 project grant to JG), the Roudnitska Foundation in France (2018-2021 research fellowship to JP); GIRACT in Switzerland (research fellowship to JP), the National Natural Science Foundation of China (grants 32070996 and 31771155 to YY), and the Science and Technology Commission of Shanghai Municipality (grant 21140900600 to YY).

