## Supporting Information for "Extracellular loop 2 of G protein-coupled olfactory receptors is critical for odorant recognition"

**Table S1.** Structure, hydrophobicity and potency of mOR256-3-wt ligands.

| Odorant | Structure | LogP*^a^* | EC50 (µM)  mOR256-3 | EC50 (µM)  mOR256-8 |
| --- | --- | --- | --- | --- |
| R-carvone | 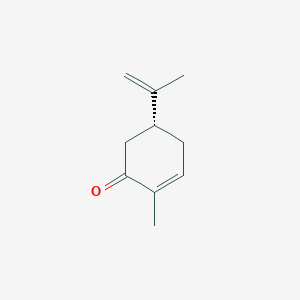 | 2.7 | 8.3 | n.a. |
| Coumarin | 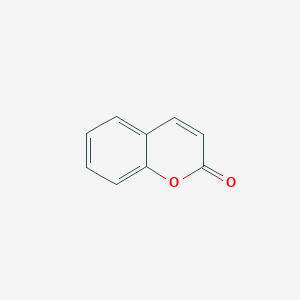 | 2.4 | 49.9 | n.a. |
| 1-octanol | 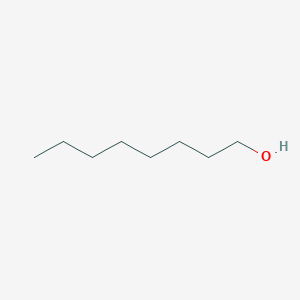 | 3.0 | 19.7 | 260.2 |
| Octanal | 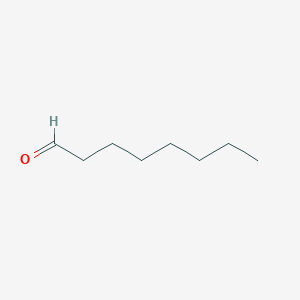 | 3.5 | 76.8 | n.a. |
| Octanoic acid | 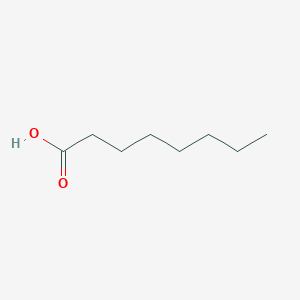 | 3.1 | 574.6 | n.a. |
| Allyl phenylacetate | 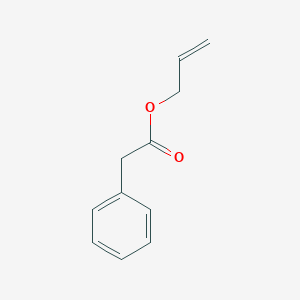 | ~2.4*^a^* | 1.4 M | n.a. |
| Benzyl acetate | 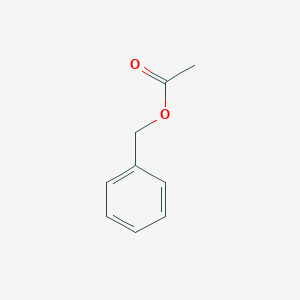 | 2.0 | 62.6 | n.a. |
| 2-heptanone | 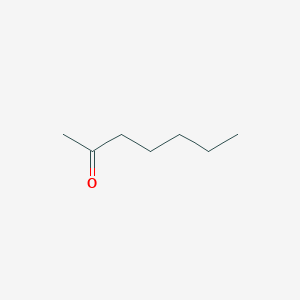 | 2.0 | 17.4 | n.a. |
| Citral | 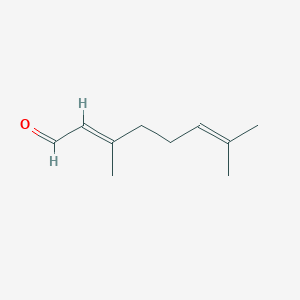 | 3.5 | 3.3 | n.a. |
| Geraniol | 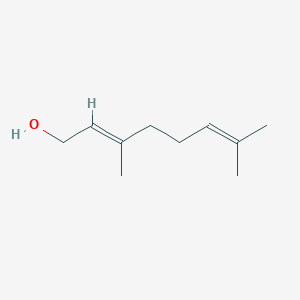 | 3.6 | 6.0 | 4.9 |

*^a^* data from PubChem

*^b^* computed by XLogP3

**Table S2.** Decoy ligands for docking

| Odorant | PubChem CID | Odorant | PubChem CID |
| --- | --- | --- | --- |
| R-(+)-Pulegone | 442495 | d-limonene | 440917 |
| 1-Butaol | 263 | Ethaol | 702 |
| 2,3-Hexanedione | 19707 | Ethyl acetate | 8857 |
| 2,5-Dimethylpyrazine | 31252 | Eugeol | 3314 |
| 2,5-Dimethylpyrrole | 12265 | Furfural | 7362 |
| Ambrette | 6753 | Hexyl octaoate | 14228 |
| 2-Methoxy-4-methylpheol | 7144 | Isobutylamine | 6558 |
| 2-Octaone | 8093 | Isobutyraldehyde | 6561 |
| 3-Methyl-2-butaol | 11732 | Isobutyric acid | 6590 |
| Acetaldehyde | 177 | Lilial | 228987 |
| Acetopheone | 7410 | Linalool | 6549 |
| Allyl hexaoate | 31266 | lyral | 91604 |
| Ammonium hydroxide | 14923 | m-Cresol | 342 |
| Amyl butyrate | 10890 | Musk ketone | 6669 |
| Amyl laurate | 62571 | Propyl acetate | 7997 |
| α-Phellandrene | 7460 | Pyridine | 1049 |
| Benzyl alcohol | 244 | Pyrrolidine | 31268 |
| Benzyl salicylate | 8363 | Thymol | 6989 |
| Phenethylamine | 1001 | Toluene | 1140 |
| Cyclohexylamine | 7965 | Cinnamaldehyde | 637511 |
| Diacetyl | 650 | Triethylamine | 8471 |
| Diethyl sebacate | 8049 |  |  |

**Table S3.** Ten top-ranked ligands from virtual screening

| Name | PubChem CID | EC50 (µM) |
| --- | --- | --- |
| Cyclohexanone | 7967 | 1.9 |
| (-)-β-Citronellol | 7793 | 2.9 |
| 2-Coumaranone | 68382 | 0.7 M |
| Ethyl vanillin | 8467 | n.s. |
| 2,4-DNT | 8461 | n.s. |
| 2-Ethyl fenchol | 106997 | n.s. |
| Acetophenone-D3 | 140244 | 4.2 |
| 4-chromanone | 68110 | 59.5 |
| 2-Nitrotoluene | 6944 | 3.6 |
| Benzaldehyde | 240 | Partial antagonist |

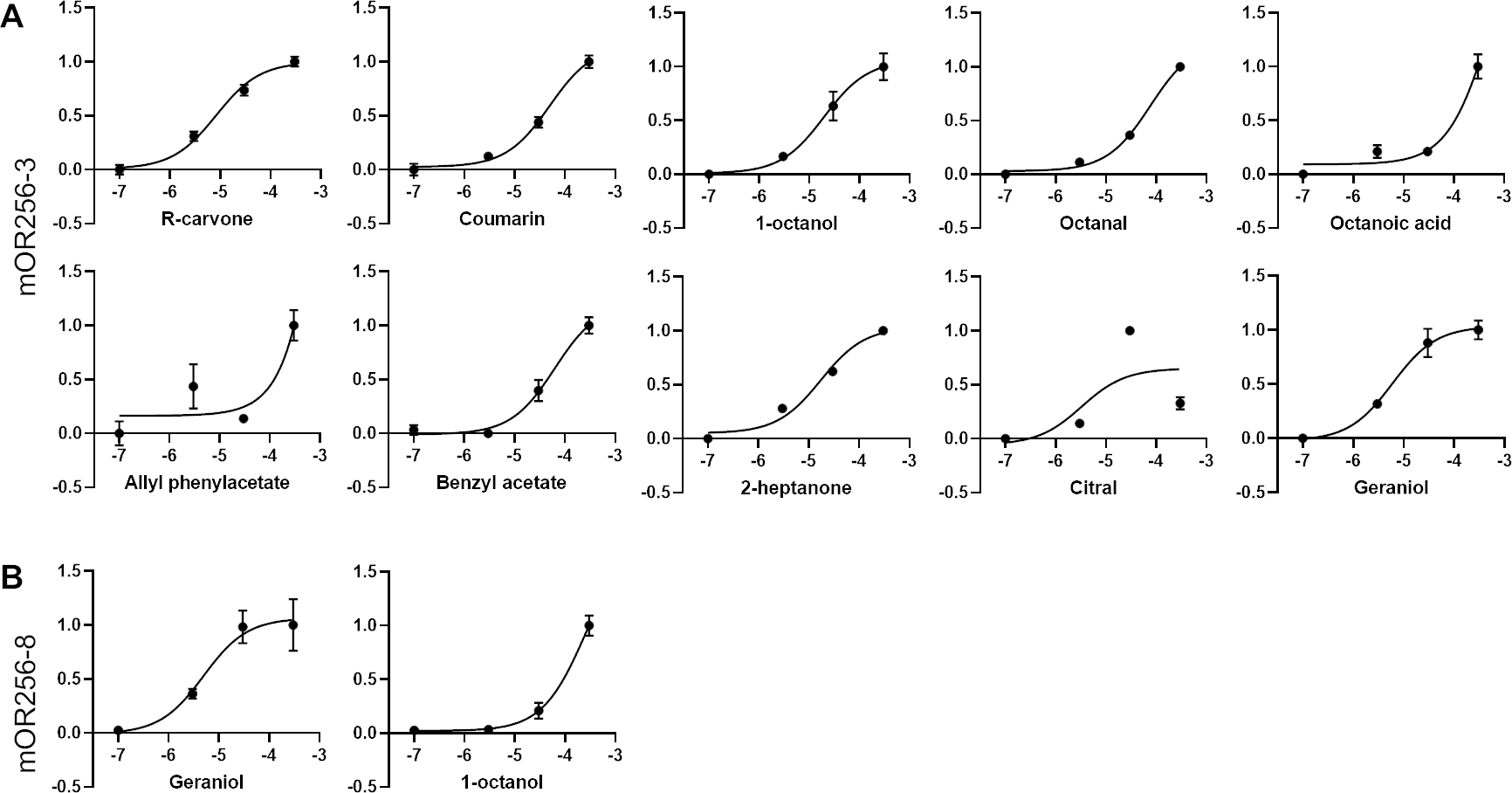

**Figure S1.** Dose-dependent response curves of **(A)** mOR256-3 and **(B)** mOR256-8 to their ligands.

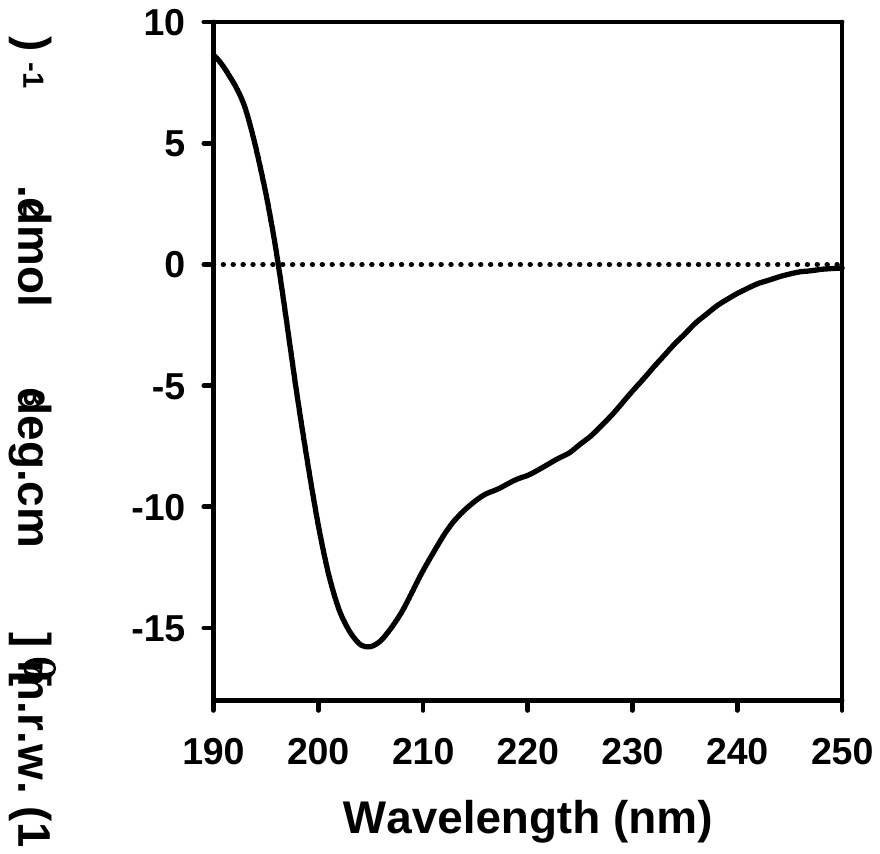

**Figure S2.** Characterization of truncated mOR256-3 ECL2 peptide using circular dichroism spectroscopy. Peptide concentration in 10 mM K_2_HPO_4_ pH 5.5, 50% TFE was 1 mg/mL. Light path: 0.01 cm.

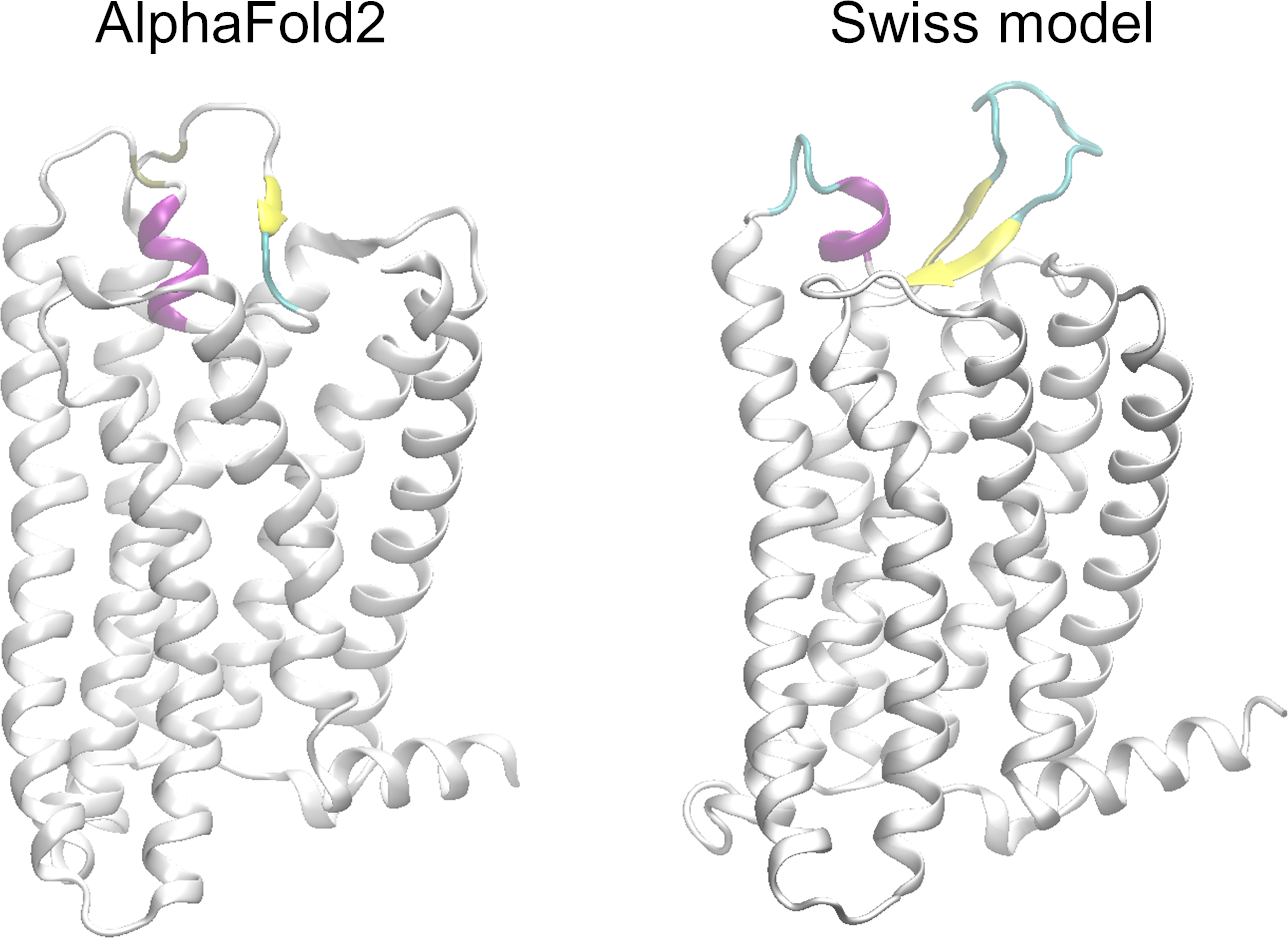

**Figure S3.** mOR256-3 models built by AlphaFold2 and Swiss model. ECL2 is colored by secondary structures. The N- and C-termini are neglected.

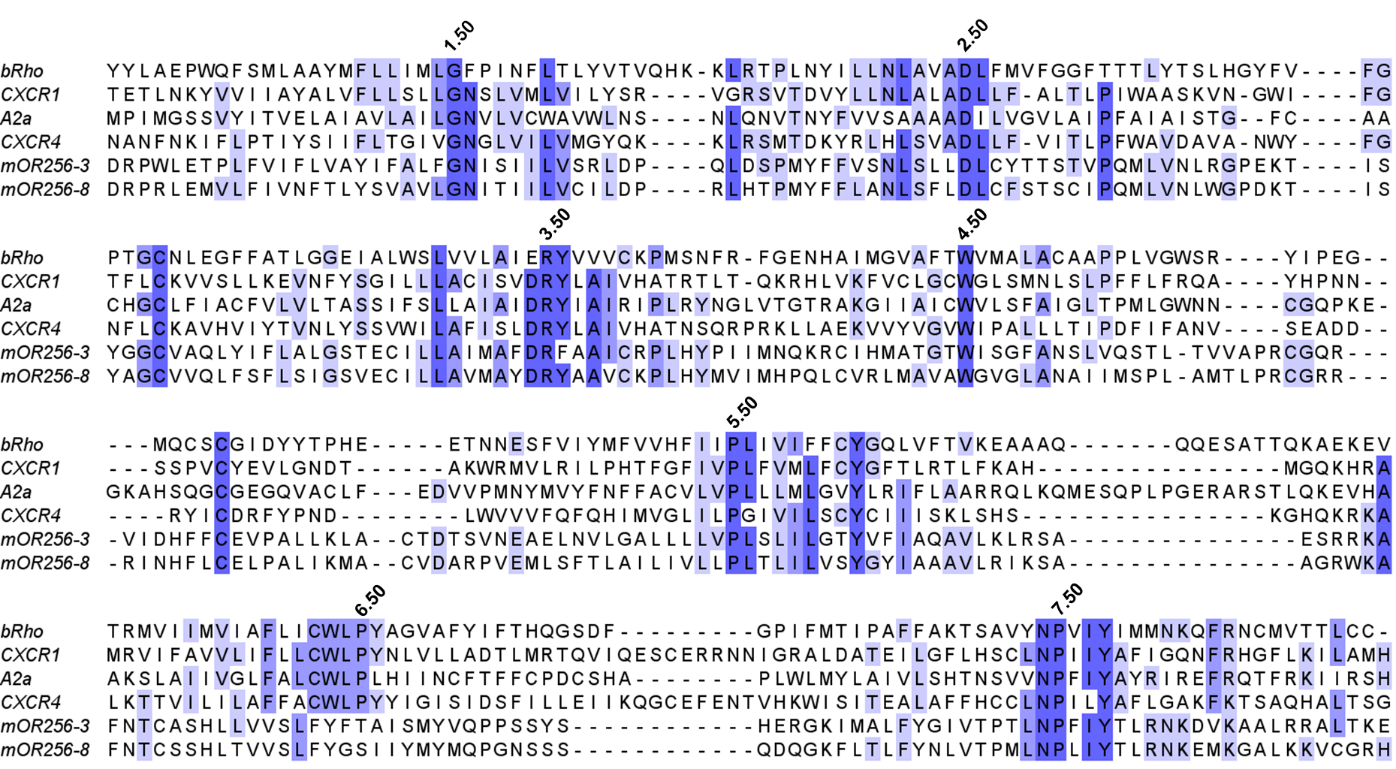

**Figure S4.** Sequence alignment for the homology modeling of mOR356-3 and mOR256-8.

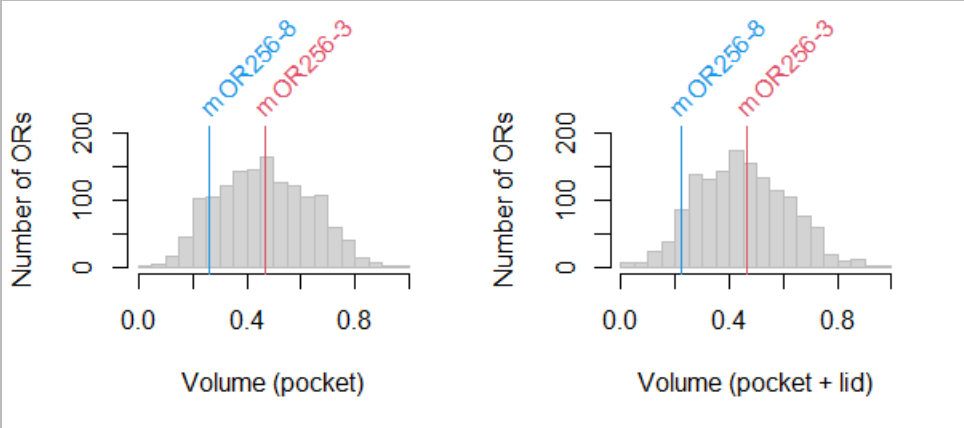

**Figure S5.** Histogram of normalized pocket volume of human and mouse ORs. The pocket volume was roughly estimated by summing up the side-chain volume of the pocket residues according to our sequence alignment and homology models.

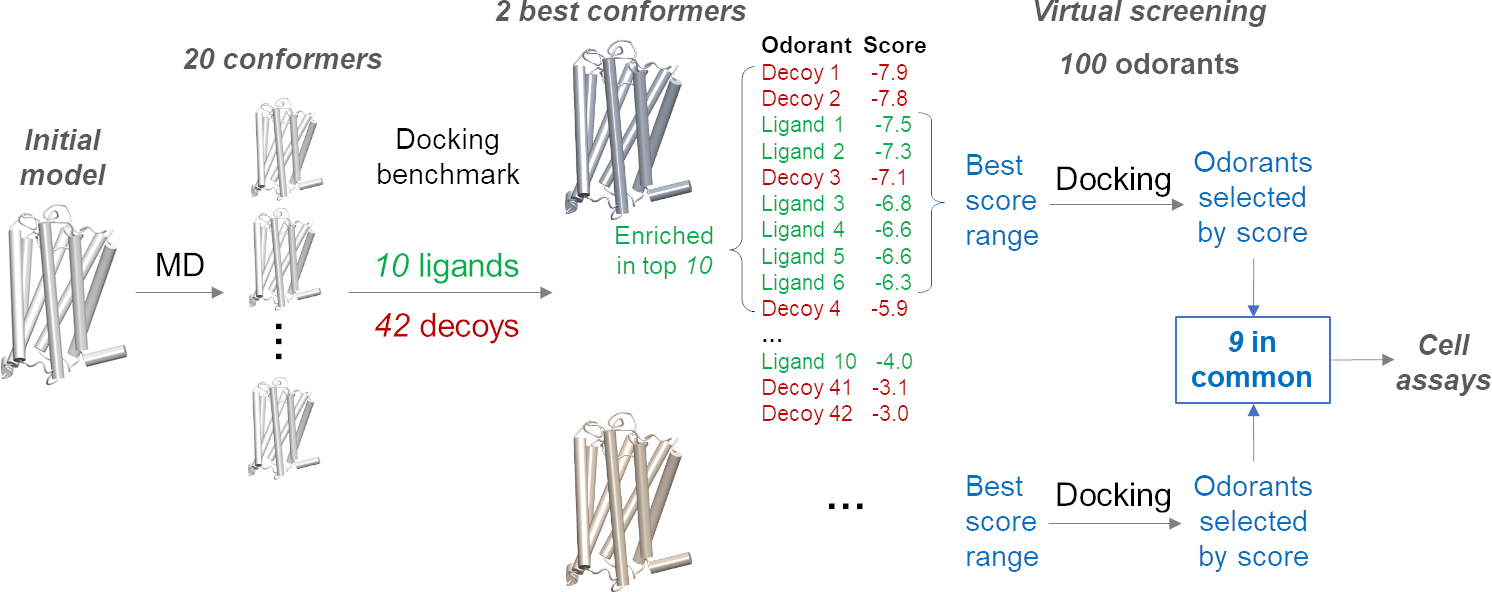

**Figure S6.** Virtual screening protocol.
